# Meta-Control Demands at the Beginning of Extended Tasks are Categorically Different from Control Demands During Their Subsequent Execution

**DOI:** 10.1101/2024.05.23.595496

**Authors:** Berhan F. Akgur, Tamer Gezici, Elif Oymagil, Ausaf A. Farooqui

## Abstract

The well-known activation of frontoparietal regions forms the basis of neuroscientific accounts of cognitive control. Control interventions that activate frontoparietal regions also generate a characteristic psychophysiological signature that, amongst others, is accompanied by increased pupil size. Here, we characterize a distinct but key aspect of cognitive control that deactivates these frontoparietal regions and decreases pupil size.

Our goals are completed through extended tasks that are hierarchically controlled and executed as a single entity despite consisting of numerous steps (e.g., ‘preparing coffee’, ‘writing emails’). This is achieved through higher-level programs instantiated at the beginning of their execution that will go on to organize cognition and instantiate various control interventions during the ensuing task execution.

Difficult episodes begin with the instantiation of more complex higher-level programs that will go on to organize a more complex set of control interventions. However, the demands related to instantiating these more complex programs are likely to be distinct from those related to control interventions. Control interventions, like selective attention, inhibition, etc., involve making goal-directed changes in sensory and motor regions to select the correct incoming information and action. In contrast, instantiating higher-level programs involves embodying the set of commands that will go on to make the relevant goal-directed neurocognitive changes later in the ensuing extended task by organizing and instantiating the correct set of control interventions.

Across four experiments, we show that while executing difficult episodes activates frontoparietal regions and increases pupil size, the beginning of these difficult episodes, i.e., junctures where more complex higher-level programs are instantiated, deactivates these same frontoparietal regions and decreases pupil size. We thus show that cognitive control related to difficult extended tasks involves categorically different demands at the beginning compared to during their subsequent execution.

## Introduction

Neurocognitive accounts of goal-directed behavior have typically focused on control interventions like attention, working memory (WM), response inhibition, rule-switching, etc. (Aron, 2007; Botvinick et al., 2004; Corbetta & Shulman, 2002; D’Esposito & Postle, 2015; Duncan & Owen, 2000; Fox et al., 2005; Niendam et al., 2012; Posner & Rothbart, 2007; Ptak, 2012). These are cognitive processes instantiated during tasks through which task-related processing is maintained in sensory and motor regions, irrelevant sensations are filtered, correct stimulus-response configurations are created, relevant representations in WM are maintained or updated, correct motor acts are selected, etc. (Aron, 2007; Desimone & Duncan, 1995; D’Esposito & Postle, 2015; Gregoriou et al., 2009; Kastner & Ungerleider, 2000; E. K. Miller & Cohen, 2001). A foundational finding in this regard has been that tasks that require more complex control interventions show greater activation of a set of frontal and parietal regions variously referred to as multiple demand (MD), attentional, or cognitive control networks (Assem et al., 2020; Duncan, 2001, 2010; Farooqui et al., 2012; Fedorenko et al., 2013; Niendam et al., 2012; Yeo et al., 2016). These control interventions are typically accompanied by an activation of modulatory cortical projections coming from brainstem regions, which correlate, amongst other psychophysiological markers, with increased pupil size (Kuipers et al., 2017; Makowski et al., 2020; Pourtois et al., 2020). Increased attention, WM load, rule-switching, response conflict, and perceptual complexity – all increase pupil size (Clewett et al., 2020; Joshi et al., 2016; Murphy et al., 2014; Porter et al., 2007; Sara & Bouret, 2012; Unsworth & Robison, 2018; van der Wel & van Steenbergen, 2018). Projections from key MD regions – frontal eye fields, intraparietal sulcus, and anterior cingulate – to pretectal olivary nucleus, superior colliculi, and locus coeruleus have been hypothesized to cause this pupil dilation (Joshi & Gold, 2020; Strauch et al. 2022). Control-related MD activations are accompanied by the deactivation of a separate set of regions referred to as the default mode regions (DMN), which has been interpreted variously as the deactivation of regions involved in internal cognition, self-processing, or mind-wandering due to resources being directed to the current task execution (Andrews-Hanna et al., 2014; Mason et al., 2007; Raichle et al., 2001). A feature of our cognition is that all our goals are achieved through extended task episodes that are extended in time and consist of numerous sequentially organized components, e.g., preparing breakfast, writing emails, executing a block of trials, etc. Control interventions that have been the subject of existing studies (e.g. attention, WM) always occur in the context of such extended episodes, which, despite being temporally extended and consisting of numerous steps, are controlled and controlled as one entity in relation to the goal (Miller, Galanter & Pribram, 1960; Schank & Abelson, 1977; Shallice & Cooper, 2011). This means that the various control interventions made during them must be instantiated as part of a larger coordinated set of goal-directed changes being made across the episode duration. A 40 s long episode, e.g., may require attention that is sustained for this duration; some junctures may require attending to parts of the environment, others may require preparing a motor act, still others may require recalling a cognitive routine, etc. Furthermore, this attention being sustained may be in relation to something being maintained in WM and, at various junctures, may be guided through the coordinated recollection of numerous episodic memories and procedures (Chen & Hutchinson, 2019; Chun & Turk-Browne, 2007; Logan, 2002; Oberauer, 2019; Taatgen, 2013). The numerous goal-directed changes made across the duration of such episodes cannot be brought about by separate and independent cognitive acts (e.g.) every millisecond, and can only be brought about via a common program that subsumes task execution, creates and evolves the cognitive focus for relevant control operations, and organizes, instantiates, and maintains various control interventions (Farooqui et al., 2023; Farooqui & Manly, 2019). Such programs would be the means through which goals retroactively organize and control the episode of cognition culminating in them (Gollwitzer & Sheeran, 2006; Greenwald, 1972; James, 1890; Jeannerod, 1988; Kruglanski & Kopetz, 2009; Lewin, 1926). These programs may thus be *metacontrol* in nature in that they are not control processes but are the means of creating the relevant organizational or set changes in cognition across the task duration, as well as the means of instantiating, coordinating, and maintaining various control processes.

Task executions begin with the instantiation of these programs (Cooper & Shallice, 2000; Farooqui & Manly, 2018b; Ruh et al., 2010; Schneider & Logan, 2006). This generates the famous finding that beginning any task episode requires additional time, which correlates with the length and complexity of the episode. Thus, when motor sequences (e.g., finger taps, button presses for piano pieces, etc.) are executed, words/sentences are articulated, memorized lists are recalled, task lists are executed, even periods of activity corresponding to an uncertain number of unpredictable trials construed as one episode executed – the first step takes the longest to initiate, and this is longer for longer/complex episodes, suggesting that the program instantiated at the beginning contains elements related to the entirety of the ensuing task episode (Anderson & Matessa, 1997; Farooqui & Manly, 2019; Henry & Rogers, 1960; Kahana & Jacobs, 2000; Klapp et al., 1973; Logan, 2004; Rosenbaum et al., 1983; Schneider & Logan, 2006). At task episode completion, these programs are dismantled. This elicits additional activity whose magnitude and spread correlate with the completed episode’s hierarchical level (Farooqui et al., 2012; Fujii & Graybiel, 2003; Sridharan et al., 2008; Zacks et al., 2001).

The higher-level and subsuming nature of these programs is also evidenced by the effect of their dismantling at episode boundaries on the switch cost related to the component steps’ rules. When the rule used to select the correct action changes, the execution efficiency decreases, and RTs and error rates increase. However, when this rule change occurs between consecutive acts that lie across the boundary of a larger task episode, this switch-cost disappears. This is because switch cost is related to a change in rule-related sets or associations between the preceding and the current acts (Rogers & Monsell, 1995; Wylie & Allport, 2000). When acts are executed as parts of larger episodes, their related sets and associations are nestled under the overarching episode-related program. Refreshing of programs at episode boundaries also dismantles any underlying rule-related sets and associations related to the component acts, leaving no disadvantage for switching rules (Farooqui & Manly, 2018b; Lien & Ruthruff, 2004; Schneider & Logan, 2015).

If these programs are the means of instantiating goal-directed control interventions across the task duration, then difficult tasks that require a more complex set of control interventions will also require a more complex program. A task episode, e.g., that requires a higher WM load and more complex updating, will also require a more complex program to organize and control these WM interventions. Such programs will take longer to instantiate, and as is well-evidenced, such is indeed the case. However, instantiating these programs is likely to be different, neurocognitively, compared to instantiating control interventions.

Control interventions are about executing a particular act and, hence, about selecting, maintaining, or updating the correct sensory or motor representations/processes or those linking the two. In contrast, instantiating these programs is about assembling the set of commands that will go on to organize a higher-level cognitive episode and instantiate relevant control interventions *later* in the ensuing task episode. Hence, the increased step 1 RT at the beginning of episodes is not driven by control demands related to the execution of any first step but by the complexity of the program related to the episode as a whole. High step 1 RT and other signs of program instantiation are seen even when what is exactly to be done in the ensuing episode is not known (Farooqui & Manly, 2018b). Likewise, episodes that have the same number of trials but last longer due to higher inter-trial intervals have higher step 1 RT. Again, this suggests that the program complexity is about the length of the ensuing cognition to be organized and executed as one entity.

We previously found that the beginning of task episodes and, hence, the instantiation of episode-related programs deactivates widespread regions, including MD regions, and the intensity of this deactivation correlates with the expected load of this program (Farooqui & Manly, 2018a). Longer tasks begin with a larger program, and this deactivation at the beginning tends to be greater at the beginning of long compared to short task episodes that are otherwise identical in their control demands (Farooqui & Manly, 2018a). The reason for this deactivation is unclear, and the following is a speculation. During task execution, MD and other regions’ activity patterns continuously move through a sequence of states, with different phases of the task corresponding to a unique neural state and, by extension, to a unique control state (Kadohisa et al., 2020, 2023; Marcos et al., 2019; Nobre & van Ede, 2023; Sigala et al., 2008; Stokes et al., 2013). This flow of neural states across the task duration requires configuring the relevant synapses at the beginning to ensure that the ensuing trajectory of neural activity patterns follows the one most optimal for goal completion. The instantiation of programs at the beginning of task episodes may correspond to this configuring of synaptic changes. This will also cease the ongoing neural activities in these regions, causing a net decrease in metabolic demands and, consequently, in BOLD activity.

In experiments 1 to 3, we built on the suggestion of our previous study (Farooqui & Manly, 2018a) and investigated if the instantiation of complex programs at the beginning of difficult task episodes is categorically different from the subsequent instantiation of control interventions during these same episodes and deactivates MD regions. Using task episodes differing in structure and content, we tested if the beginning of difficult episodes that require a more complex program deactivates the same MD regions that will subsequently activate during the execution of these difficult episodes.

Experiment 4 investigated this issue using pupil size. Any large-scale change in cortical activity is accompanied by changes in the modulatory brainstem systems that project diffusely to various cortical regions (Reimer et al., 2014). The activity of many of these systems (popularly adrenergic but also cholinergic, serotonergic, and even hypocretinergic) correlates with pupil size (Joshi et al., 2016). Control interventions and MD activations are typically accompanied by an increased pupil size (Mathôt, 2018; Strauch et al., 2022; van der Wel & van Steenbergen, 2018). If program instantiation at the beginning of difficult episodes deactivates MD regions, it may also cause a decrease in pupil size. We tested this in Experiment 4.

## Results

In experiments 1 and 2, we used a standard block design with easy and hard task episodes, each lasting around 15-30 s. Contrasting hard versus easy episodes is well known to show increased activation of MD regions. We investigated if the beginnings of hard episodes, when more complex overarching programs get instantiated, deactivate MD regions more than the beginnings of easy episodes.

### Experiment 1

Participants executed hard and easy episodes of an auditory n-back task that involved maintaining and updating either 3 (hard episodes) or 1 (easy episodes) item (Figure 1a). An episode consisted of 10 trials. On each trial, a letter was presented. For the 1-back task blocks, participants were to keep the first trial letter in mind and proceed to the next trial by pressing a button. From the second trial onwards, they were to respond if the letter they heard was the same as the previous trial (index finger: yes; middle finger: no). For the 3-back task blocks, the trial content was the same, but participants now had to remember the letters of the first three trials, and from the fourth trial onwards, they had to decide if the current letter was the same as the one they heard three trials earlier. Easy and hard blocks alternated with each other, and their order was counterbalanced across participants. Participants executed a total of 10 easy and 10 hard blocks. Participants were expectedly slower and more erroneous during 3-back compared to 1-back episodes (Figure 1; F_1,20_ = 56.682, p < 0.001).

**Figure 1.**
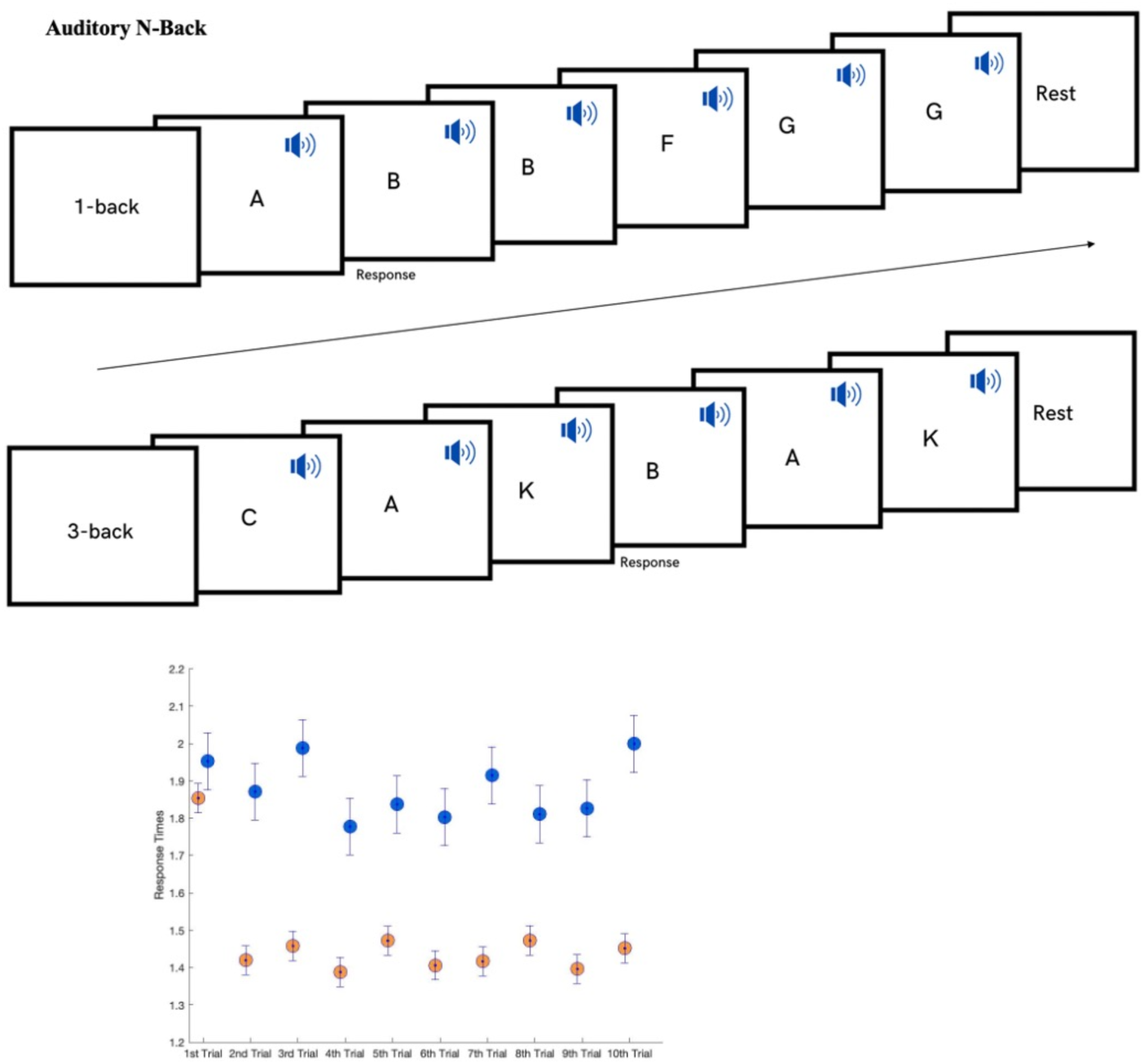
Experiment 1. Auditory working-memory updating: Each episode had 10 trials. Participants decided if the letter they heard on a given trial was the same as that on 1 (on easy blocks) or 3 (on hard blocks) trials prior (see Methods). Behaviorally, RTs were higher on hard (blue) compared to easy (orange) blocks.

Since the 10-trial block is presented as a single task episode that starts with trial 1 and ends with trial 10, it will likely be executed via a single program instantiated at the beginning. Hard episodes will require a more demanding program at their beginning that will subsequently instantiate a more complex set of control interventions. To look at the neural correlate of instantiating a more demanding program, we contrasted the phasic activity elicited at the beginning of hard episodes against that elicited at the beginning of easy episodes. This phasic activity at the beginning was captured by an event of no duration, modelling the beginning of these episodes. If program instantiation deactivates widespread regions including MD regions, then instantiating more demanding programs at the beginning of hard episodes should further deactivate these regions.

In contrast to this phasic activity at the beginning, the sustained activity elicited across the duration of these episodes, captured by modeling the episode duration as an epoch, is related to the execution of these episodes. The difference in this activity between hard and easy episodes reflects the more complex control interventions made during the hard episodes, which are well-known for activating MD regions. We expected the same here. We can, thus, predict that the difference in MD activity during easy and hard episodes will be different at their beginning compared to during their subsequent execution.

We first show unthresholded t-maps so that regions that show trends that do not reach statistical significance also get displayed. Figure 2 shows regions where activity at the beginning and during the execution of hard episodes was higher (hot colors) and lower (cold colors) than the corresponding parts of easy episodes. The beginning of hard episodes resulted in a decrease in activity in widespread brain regions, including MD regions, compared to that of easy episodes. In contrast, the activity during the execution of the hard episode was higher in MD regions.

**Figure 2.**
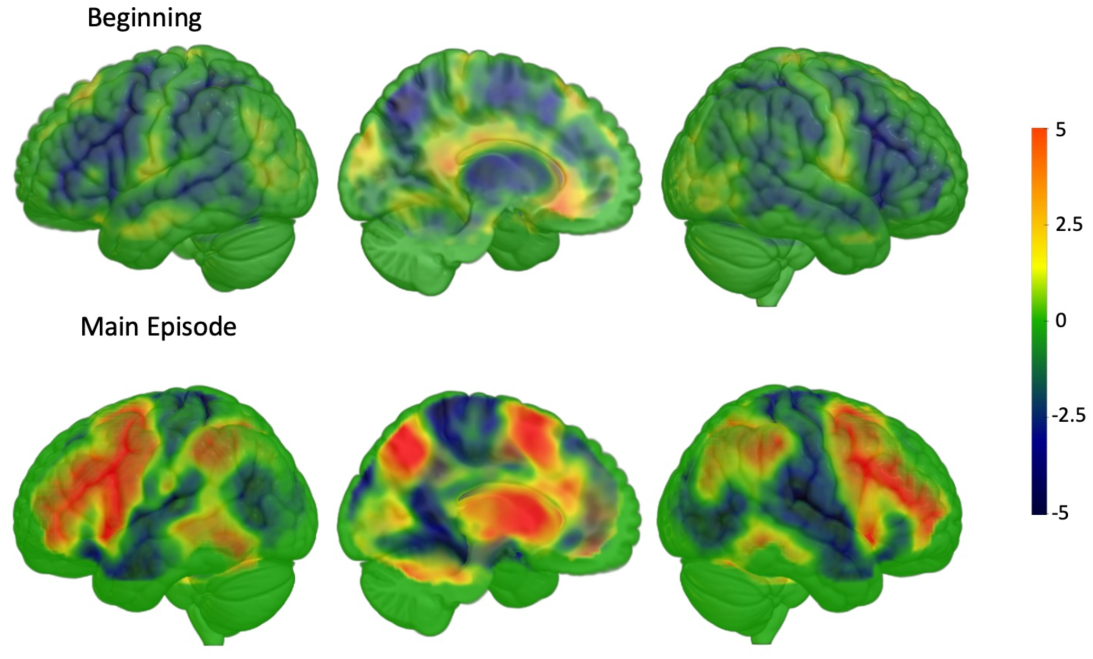
Experiment 1. T-maps of the contrast hard>easy at the beginning and during the main episode. MD regions activated more for hard episodes only during the main episode epoch. At the beginning of the episode, MD regions, along with other regions, decreased their activity for hard episodes. The t threshold corresponding to an FDR correction at p < 0.05 is at 2.9 and 2.4 for the top and bottom images.

A whole brain contrast (Figure 3) looking for this differential response to difficulty at the beginning and during the subsequent execution of the episode showed significance at all MD regions – bilateral inferior frontal sulcus extending up to the anterior prefrontal cortex, anterior insula, intraparietal sulcus, and pre-supplementary motor area. As evident in the plots of these regions, in all of them, the beginning of hard episodes led to a stronger deactivation, but their subsequent execution led to a stronger activation. Thus, the demands of executing difficult episodes were distinct at their beginnings when they required a complex meta-control program compared to during their subsequent execution when they required a complex set of control interventions.

**Figure 3.**
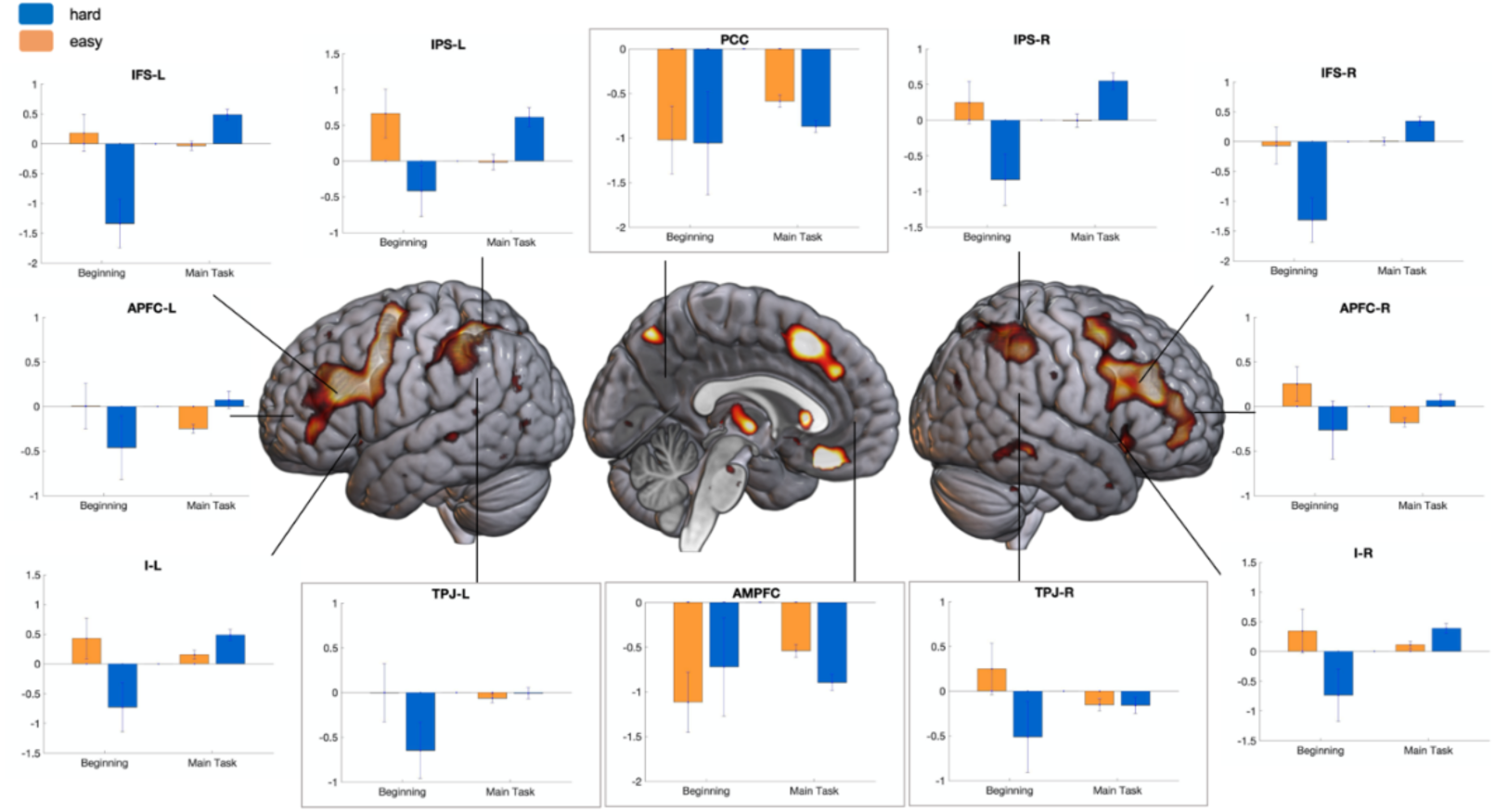
Experiment 1. Brain render shows regions where the effect of difficulty was different at the beginning compared to during the main episode (FDR, p < 0.05). Plots show the estimates of activity at the beginning and during the task episode for easy and hard conditions in MD regions showing significance in the whole brain contrast. In all of them, the beginning of hard episodes elicited lower activity than the beginning of easy episodes; however, these same regions activated more during the subsequent execution of these hard episodes. Error bars represent 95% confidence intervals. Plots in the inset show DMN regions. Note the difference between temporoparietal junctions’ (TPJ) response patterns from those of the posterior cingulate (PCC) and anteromedial prefrontal cortex (AMPFC). TPJ, especially on the right, showed decreased activity on hard step 1 but not on hard step 2. In contrast, PCC and AMPFC showed the opposite – decreased activity on hard step 2 but not on hard step 1 (this interaction being significant, F5,145 = 6.2, p < 10^-4^).

While we had no hypotheses about DMN regions beyond expecting a more intense deactivation during hard compared to easy episodes, we serendipitously discovered a dissociation between midline DMN regions – anteromedial prefrontal cortex (AMPFC) and posterior cingulate cortex (PCC), from those of the temporoparietal junction (TPJ). Temporo-parietal junctions, bilaterally, showed lower activity at the beginning of hard compared to easy episodes but showed no difference between these episodes during their subsequent execution. PCC and AMPFC, in contrast, showed the opposite – no difference at the beginning but a greater deactivation during the subsequent execution of hard episodes. This interaction between effects of difficulty, junctures of execution (beginning vs. during the main episode), and ROIs (bilateral TPJs vs. PCC and AMPFC) was significant (F_1,20_ = 10.2, p = 0.004).

As we describe below, these same findings were seen in experiment 2, which had a very different task content.

### Experiment 2

Participants executed hard and easy episodes involving 5 tactile judgment trials (Figure 4). On each trial, they were presented with a pair of shapes and were to find the larger shape. Shape pairs presented during hard episodes had a smaller difference in size and were more difficult to discern (see Methods). Behaviorally, participants were slower and more erroneous during hard episodes (F_1,20_ = 2603, p < 0.001; Figure 4).

**Figure 4.**
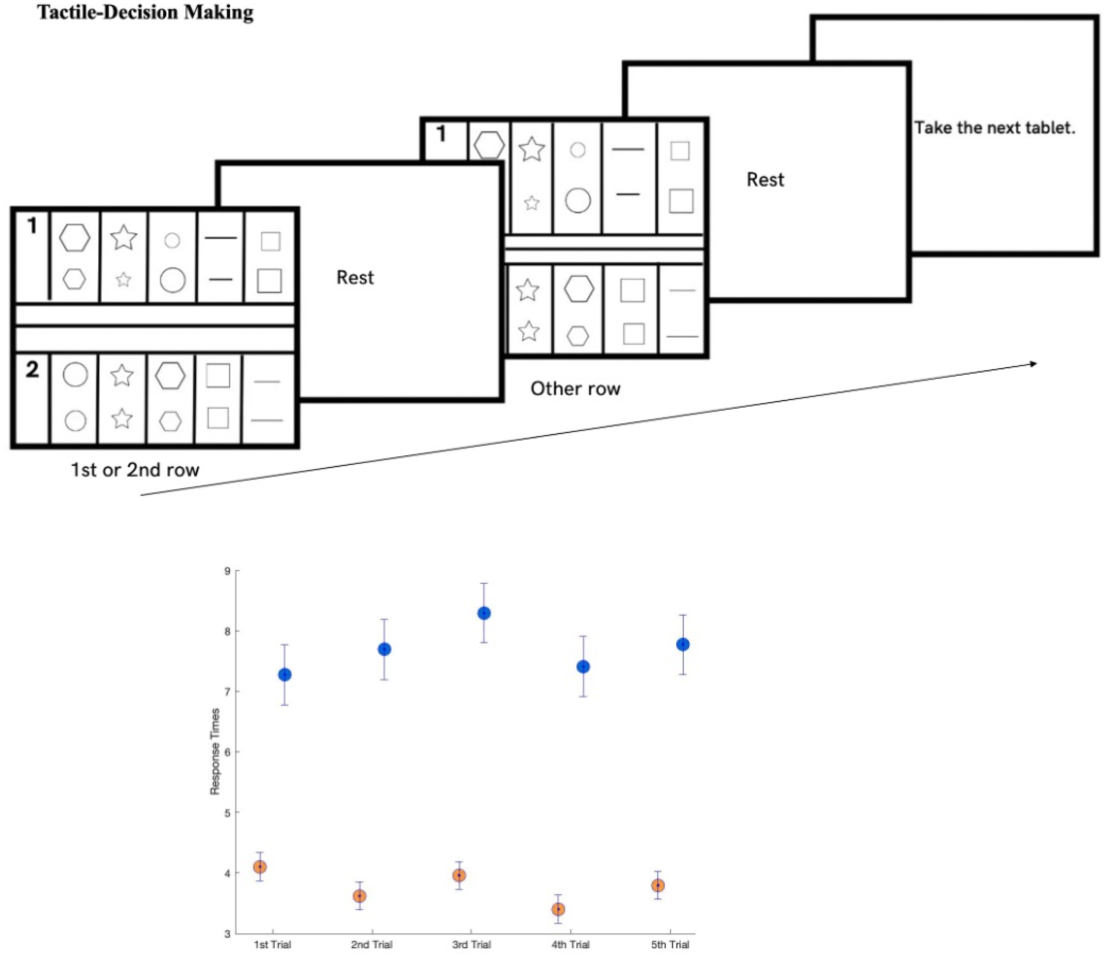
Experiment 2. Tactile Decision-Making: Participants decided which of the two shapes was larger in size. The task was done through a plexiglass tablet that had an easy and a hard part, each of which had 5 trials. Each trial involved a pair of shapes drawn with raised margins. Margins of shapes on easy trials were more raised above the surface, making them easier to perceive by touch. Further, the size difference between the shapes of a pair was larger on easy trials.

We again compared the hard and easy episodes in terms of the phasic activities elicited at their beginning (to capture the activity related to program instantiation) and sustained activity elicited during their execution (to capture activity related to the control interventions made during their execution).

Figure 5 shows regions where activity at the beginning and during the execution of hard episodes was higher (hot colors) and lower (cold colors) than the corresponding parts of easy episodes. The beginning of hard episodes again resulted in a decrease in activity in widespread brain regions, including MD regions, compared to that of easy episodes. In contrast, the activity during the subsequent execution of the hard episode was higher in MD regions. This differential effect of difficulty at the beginning versus during the execution of the episode, as shown by a whole brain ANOVA, was significant in most MD regions – inferior frontal sulcus, reaching up to the anterior prefrontal cortex, anterior insula, presupplementary motor cortex and intraparietal sulcus (Figure 6).

**Figure 5.**
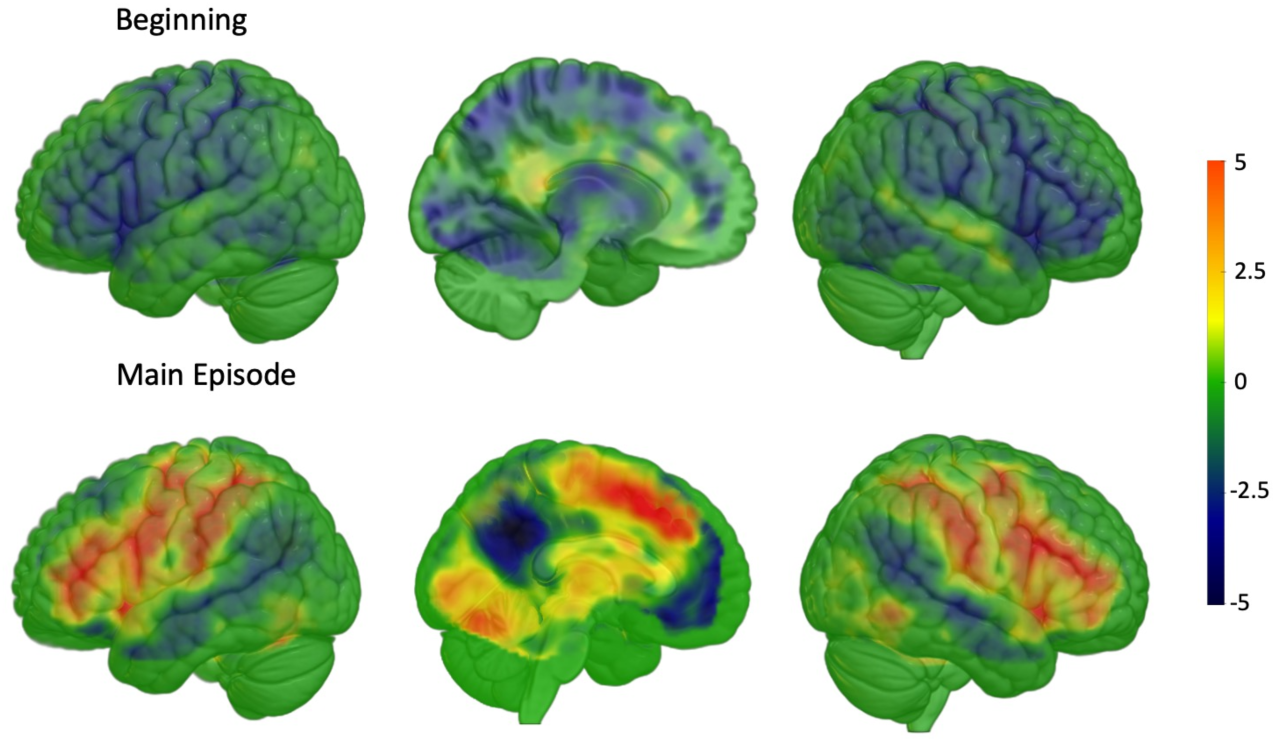
Experiment 2. T-maps of the contrast hard>easy at the beginning and during the main episode. Again, the MD regions activated more for hard episodes only during the main episode epoch. At the beginning of the episode, MD regions, along with other regions, decreased their activity for hard episodes.

**Figure 6.**
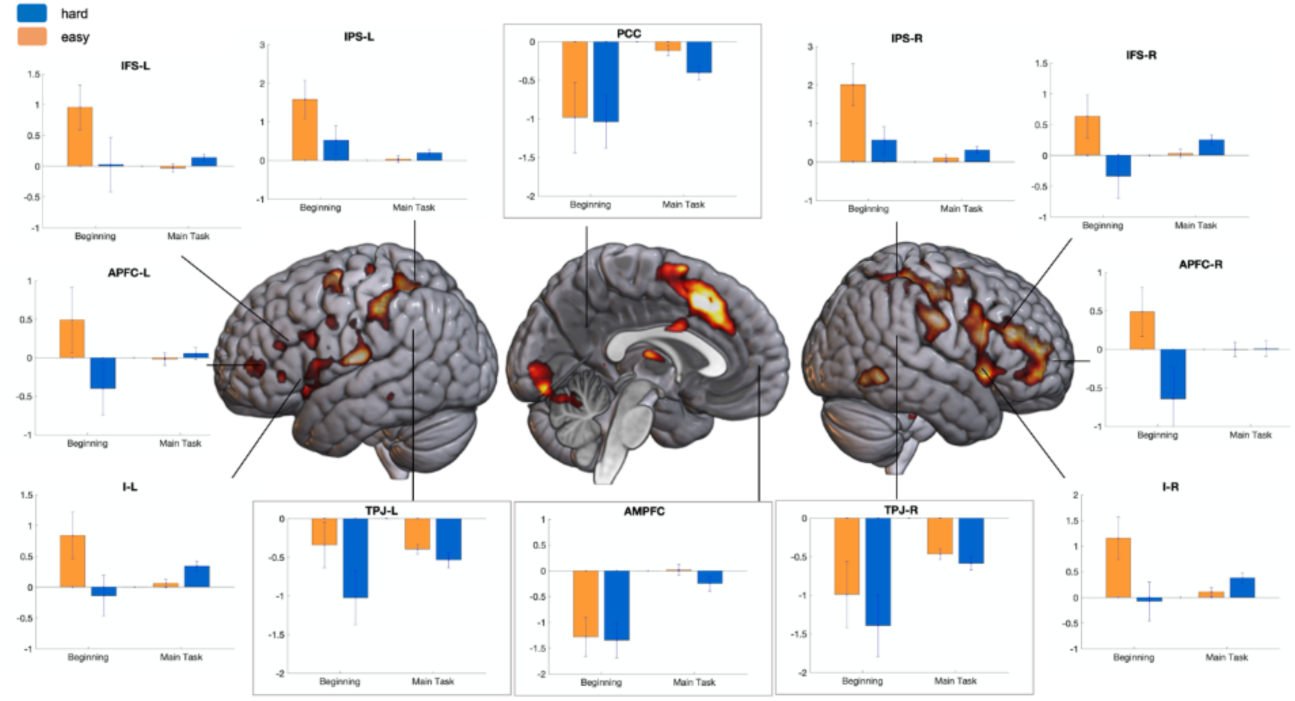
Experiment 2. Brain render shows regions where the effect of difficulty was different at the beginning compared to during the main episode (FDR, p < 0.05). Plots show the estimates of activity at the beginning and during the task episode for easy and hard conditions in MD regions showing significance in the whole brain contrast. Note the similarity with Experiment 1 (Figure 3). Again, in MD regions, the beginning of hard episodes elicited lower activity than the beginning of easy episodes; however, these same regions activated more during the subsequent execution of these hard episodes. Error bars represent 95% confidence intervals. Plots in the inset show DMN regions. The difference seen in Experiment 1 in the responses of bilateral TPJs from those of medial DMN regions – PCC and AMPFC – was also present here (F1,20 = 5.1, p = 0.03). TPJs deactivated more at the beginning of hard compared to easy episodes. PCC and AMPFC did not show this difference but deactivated more during the execution of the main episode.

The difference we had discovered in Experiment 1 between the responses of bilateral TPJs from that of midline DMN regions, PCC and AMPFC, was also present in this experiment. Bilateral TPJs, but not PCC and AMPFC, deactivated more intensely at the beginning of hard compared to easy episodes. PCC and AMPFC, in contrast, showed more intense deactivation during the hard compared to easy episodes. This interaction between effects of difficulty, junctures of execution (beginning vs. during the main episode), and ROIs (bilateral TPJs vs. PCC and AMPFC) was again significant (F_1,20_ = 5.1, p = 0.03).

### Experiment 3

Here we made a decisive test of our thesis by making the first step of difficult episodes have a higher WM demand than the first step of easy episodes. WM demand is considered as a well-known activator of MD regions. However, if the MD deactivation at the beginning of difficult episodes is a stronger effect, then step 1 of these episodes will elicit a relative deactivation despite having a higher WM demand.

The experiment design described below was also used in Experiment 4. Participants did an extended task that consisted of four steps, organized as two interleaved subtasks (Figure 7). The task began with step 1. One (in easy episodes) or two (in hard episodes) oblique lines, hence referred to as subtask-B lines, whose orientations were to be kept in mind, were presented. These stayed on till a button was pressed. Participants then saw two other lines (subtask-A lines) whose orientations were also to be kept in mind (step 2). These, too, stayed on till a button was pressed. Participants were then probed to recall the orientation of one of the subtask-A lines by orientating the probe to the same angle (step 3). Lastly, they were probed to recall the orientation of one of the subtask-B lines, presented during step 1, by orientating the probe (step 4). Easy and hard task episodes required that the orientation of one and two subtask B lines, respectively, presented in step 1, be kept in mind till the end of the episode (step 4) while being shielded from the interference of subtask-A lines.

**Figure 7.**
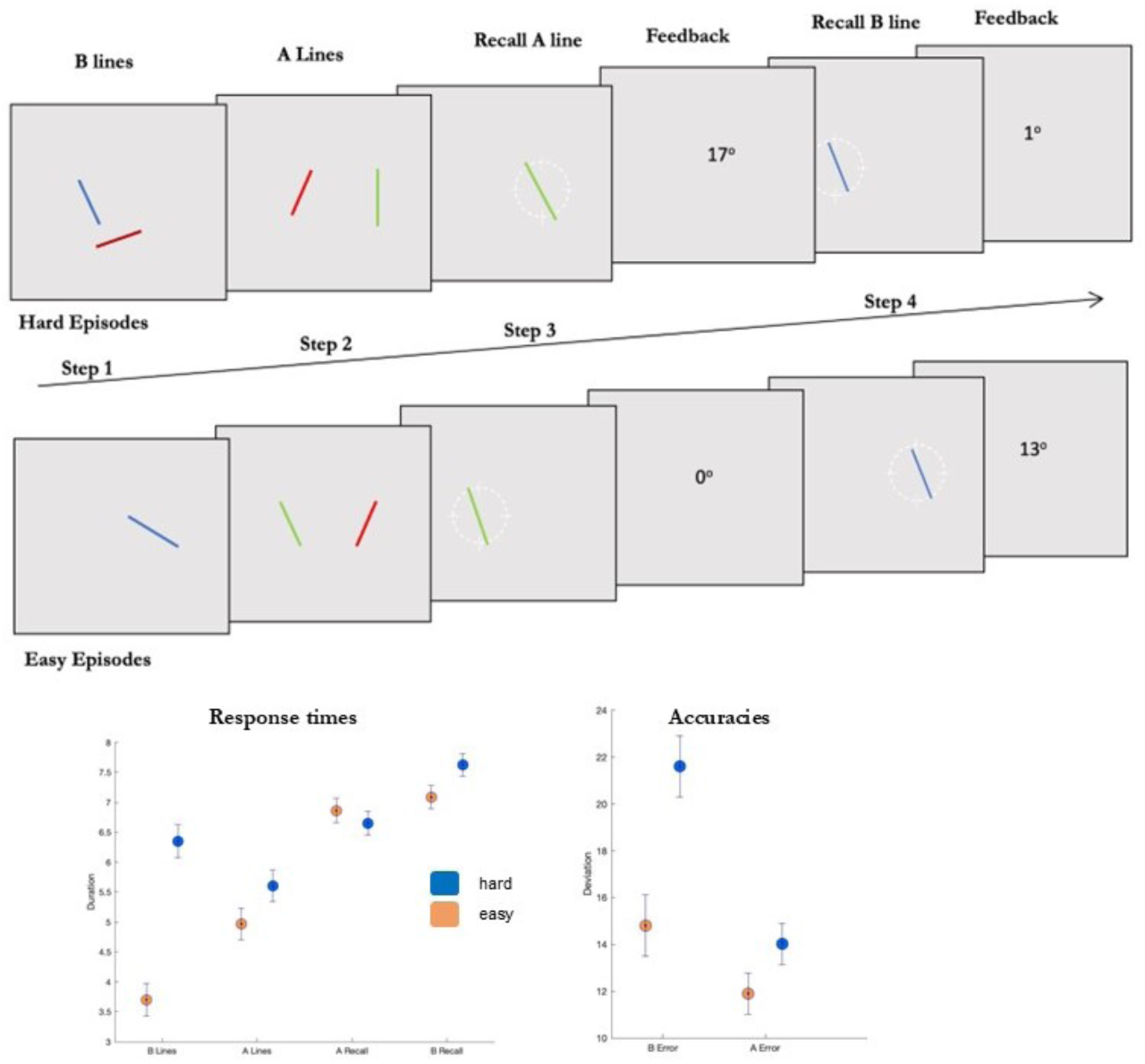
(Experiments 3 and 4). Participants executed two kinds of extended task episodes. Hard episodes began with the presentation of two lines whose orientations were to be kept in mind for subtask-B (step 1). They then (step 2) saw two different lines (referred to as subtask-A lines), kept their orientations in mind, and were probed to recall the orientation of one of these by orienting the probe to the same angle (step 3). After finishing this subtask-A, they were probed to recall the orientation of one of the subtask-B lines (step 4). Easy episodes were identical to hard episodes except that they started with a single subtask-B line presentation. Hard and easy episodes thus differed in the number of subtask-B lines that were to be kept in mind across the duration of the episode while subtask-A was being executed.

Hard episodes involved a more complex episode that required a more complex set of control operations, e.g., a more complicated set of operations to keep subtask-A lines from interfering with subtask-B lines. Step 1 of these episodes will, therefore, require a more complex episode-related program to instantiate these control operations across the episode that (as evidenced in Experiments 1 and 2) deactivates MD regions. But step 1 of these episodes also involves a higher WM load, known to activate MD regions. This step, thus, pits these two effects against each other. A deactivation of MD regions at this step would be strong evidence for our claim and cannot be explained by existing accounts for whom increased WM load should activate these regions. In comparison, by step 2, the program is already in place, and hard episodes require more complex WM operations (instantiated through this program) to keep line representations related to the two subtasks from interfering. Hence this step can only be expected to activate MD regions.

Since the design was identical for experiments 3 and 4, we combined their behavioral results. Participants expectedly took longer to register subtask-B lines at step 1 of hard compared to that of easy episodes, and they took longer to recall these lines at step 4 of these episodes (Figure 7; t_59_ > 6, p < 0.001). However, these times were not different for subtask A-lines across hard and easy episodes (t_59_ < 1.46, p > 0.15). Recollections were more erroneous during hard episodes for both subtasks B and A-line orientations (t_59_ > 3.7, p < 0.001).

The brain-render in Figure 8 shows regions whose response to difficulty was different across steps 1 and 2. These included key MD regions – bilateral inferior frontal sulci extending into the anterior prefrontal cortex, anterior insula and intraparietal sulci. As evident in the plots of activity time-series of these regions, at step 1, their activation was lower during hard compared to easy episodes; in contrast, at step 2, their activation was higher during hard episodes. Thus, even though step 1 of hard episodes involved a higher WM load, it deactivated these MD regions compared to step 1 of easy episodes. Step 2 of these hard episodes expectedly activated MD regions. A categorically different response of MD regions to difficult task episodes at steps 1 and 2 is again evidence that these positions involve categorically different issues.

**Figure 8.**
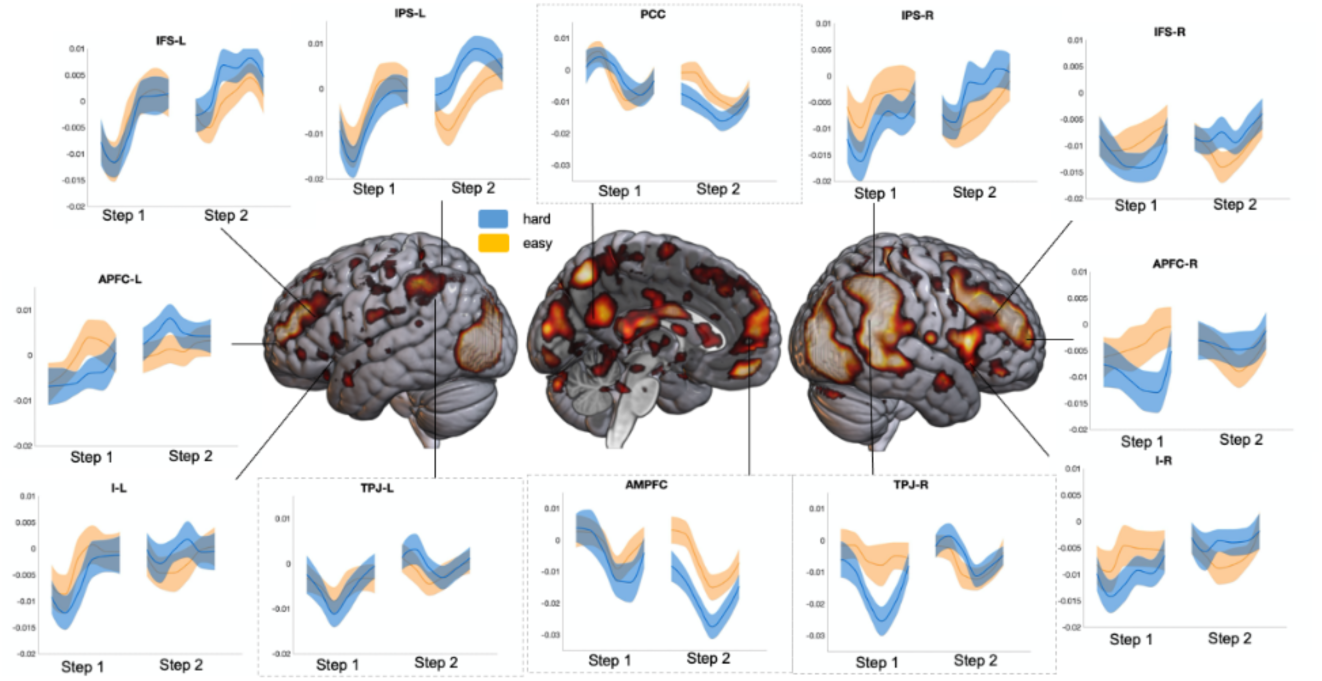
(Experiment 3). Whole-brain render shows regions where the effect of difficulty was different across steps 1 and 2 (FDR, p < 0.05). Plots show the time-series of activity across 12 seconds following the onset of steps 1 and 2. In all MD regions, hard step 1 elicited lower activity, while hard step 2 elicited higher activity than analogous easy steps (F5,145 > 2.3, p < 0.04 for all ROIs depicted). Plots in the inset show DMN regions. Note the difference in TPJ’s response patterns from that of PCC and AMPFC. TPJ, especially on the right, showed decreased activity on hard step 1 but not on hard step 2. In contrast, PCC and AMPFC showed the opposite – decreased activity on hard step 2 but not on hard step 1 (this interaction being significant, F5,145 = 6.2, p < 10^-4^).

For the third time in 3 experiments, we again noted a dissociation between midline DMN regions – anteromedial prefrontal cortex (AMPFC) and posterior cingulate cortex (PCC), from those at the temporoparietal junction (TPJ). During hard episodes, TPJs showed decreased activity at step 1 but increased activity at step 2 (F_1,29_ = 7.7, p < 0.01). In contrast, AMPFC and PCC had nearly the opposite response. These showed no difference at step 1 but decreased activity at step 2 of hard compared to easy episodes (F_1,29_ = 7.4, p = 0.01). Bilateral TPJs’ behavior significantly differed from that of PCC and AMPFC (interaction between the effects of difficulty, step, and ROIs, F_5,145_ = 6.2, p < 10^-4^).

### Experiment 4

Control interventions that activate MD regions also increase pupil size (Mathôt, 2018; Strauch et al., 2022; van der Wel & van Steenbergen, 2018). If the instantiation of episode-related programs deactivates MD regions, then it may also decrease pupil size. We used the same design as in Experiment 3. Since WM demands are also well known to increase pupil size (Kahneman, 1973; Unsworth & Robison, 2018), step 1 thus pits the effects of WM demand and the possible effects of program instantiation against each other since it involves both a higher WM demand and a more complex episode-related program instantiation. Step 2, in comparison, only involves WM demands.

As evident in Figure 9A, at step 1, pupil size decreased during hard compared to easy episodes despite the former involving a higher WM load (repeated measures ANOVA, F_1, 29_ = 8.2, p < 0.01). At step 2, in contrast, hard episodes had a larger pupil size (F_1, 29_ = 4.9, p = 0.03). Crucially, as with MD activity, the effect of difficulty at steps 1 and 2 was significantly different (F_1,29_ = 12.3, p = 0.001). Thus, paralleling MD deactivation, step 1 of hard episodes had a smaller pupil size.

**Figure 9.**
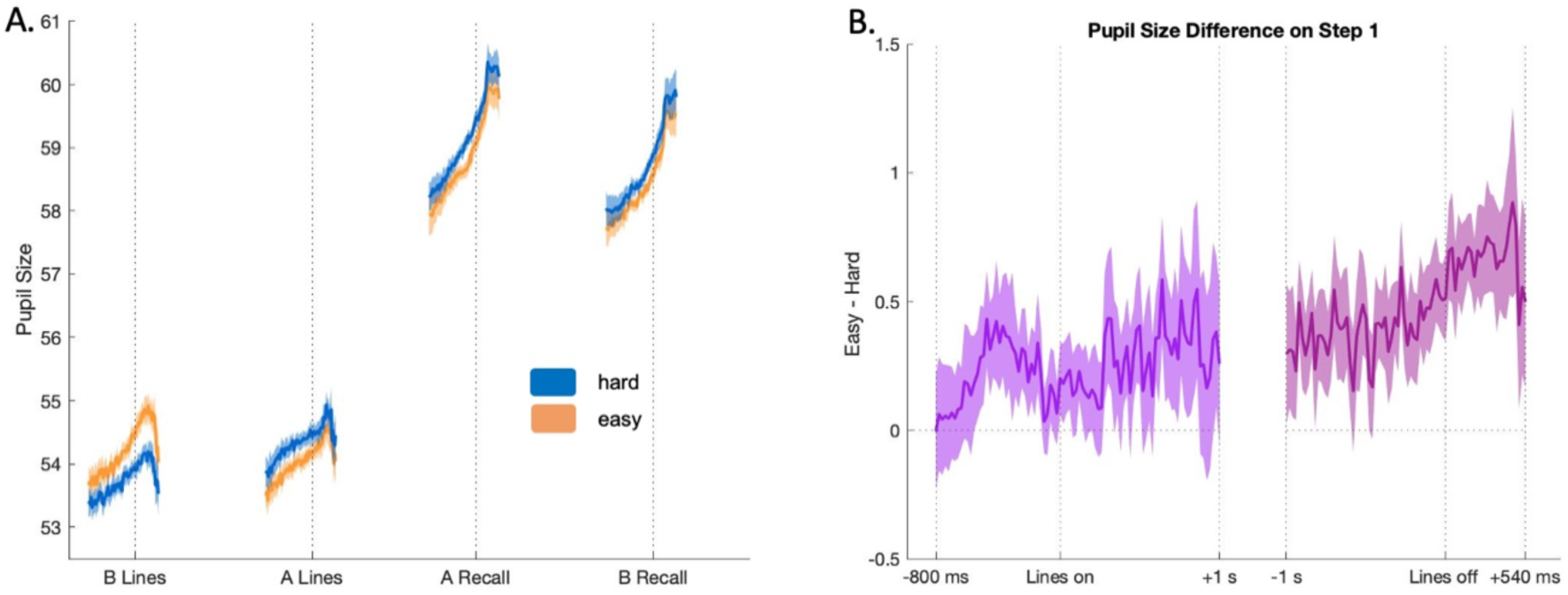
(Experiment 4). A. Time-series of pupil size across the four steps of the hard and easy task episodes. Line plots represent mean values, and shadings show 95% confidence intervals. Dotted vertical lines represent the time when participants pressed the button marking the completion of the respective step. Plots show pupil size 1s before and 500 ms after this completion. B. Time-series of pupil size difference between easy and hard step 1 (Easy – Hard) during different parts of step 1. The first plot (violet) shows the time-series of difference from 800 ms before to 1 s after the onset of subtask-B lines. Note that the difference between easy and hard step 1s increases immediately *prior* to the onset of these lines but decreases after their onset. The second plot (lilac) shows the time-series of difference from 1 s before to 540 ms after the offset of subtask-B lines. Note that the difference increases immediately *after* the offset. Both show that the pupil size difference cannot be driven by the visual differences between easy and hard step 1 because if such was the case, then the difference should have been higher while the lines were on and not prior to their onset and after their offset.

While the easy and hard screens at step 1 were equal in illumination, it can be suspected that the requirement to focus on an additional colored line during hard step 1 caused the pupil size to decrease. However, if this or any other visual issue resulting from the presence of an additional line during hard step 1 was the cause of decreased pupil size, the pupil size difference between easy and hard conditions would primarily be present while the lines were on and the difference would be absent before the onset and after the offset of these lines. However, as evident in Figure 9B, the opposite was the case. The difference between pupil sizes on step 1 of easy and hard episodes shows a higher difference (with pupil sizes being larger on easy episodes) immediately before the onset of lines on the screen and immediately after their offset. In fact, the difference was highest in the 500 ms period after the lines disappeared. The decreased pupil size on step 1 of hard episodes cannot, therefore, be due to visual issues resulting from the presence of an additional line.

## Discussion

While the activation of MD regions and increased pupil size *during* the execution of difficult task episodes is a very well-known finding, we showed that the *beginning* of difficult task episodes, when higher-level programs related to the whole episode are assembled, deactivates these same regions and decreases pupil size. These findings were identical across task episodes involving auditory WM updating, visual working memory interference, and tactile perceptual judgment, which shows that these are general features of any extended task episode regardless of its content and structure.

Crucially, these findings can only be explained in terms of the overarching, higher-level episode-related entities. Disregarding these entities, one will have to explain the deactivation and decreased pupil size in terms of the demands intrinsic to the first step. The first step in these experiments required registering a letter in auditory WM, making tactile judgments, and registering line orientations in visual WM. This requires positing something that is common across these different task contents and deactivates MD regions and then explaining why these same demands later in the episode activated these very same regions. Explaining the latter forces one to admit that there must be something unique about starting an episode that led to this response. However, it’s not enough to posit episode-initiation effects because the finding is that initiating hard episodes elicits a different neural and psychophysiological response than initiating easy episodes (i.e. widespread deactivation and decreased pupil size). This forces one to admit that the preparatory goings-on at the beginning of hard episodes are distinct from those of easy episodes. And this is what we posit, except that we use the term episode-related program for these preparatory goings-on because, in light of other findings, these preparatory goings-on create a hierarchically organized cognition (Farooqui et al., 2012, 2023; Farooqui & Manly, 2018a, 2018b; Schneider & Logan, 2006, 2015).

Control of extended task episodes is well-recognized to require a hierarchical cognition that has entities (representations/operations/processes) at multiple levels, with the higher-level entities controlling the lower-level ones (Broadbent, 1977; G. A. Miller et al., 1960; Norman & Shallice, 1986). Programs instantiated at their beginning are fundamental to this in that they are the higher-level entities and are the means of controlling and executing extended tasks as one unit (Farooqui & Manly, 2018a). However, as evident in the above results (Figures 2 and 5), instantiating more demanding programs at the beginning of hard episodes were not accompanied by increased activity in any brain region. This is very different from the expectations of many accounts of hierarchical cognition. For a set of them, more anterior regions of the PFC activate to higher-level entities (e.g. Badre & Nee, 2018; Christoff et al., 2009; Koechlin, 2003). For others, the DMN may be about the higher-levels of cognition and may represent temporally extended episodes in cognition (Margulies et al., 2016; Speer et al., 2007; Wen et al., 2020). Some have also suggested that MD, especially lateral prefrontal, regions may be the ones instantiating programs that control and execute extended sequences of thought and behavior (Duncan, 2013; Nee & D’Esposito, 2016). These accounts would, respectively, predict that anterior prefrontal regions, DMN, and MD regions will have higher activation at the beginning of more demanding task episodes. In contrast, we showed that the demands related to more complex programs at the beginning of difficult tasks deactivate widespread regions, including MD regions.

Activation and deactivation to cognitive demands are well-known but have been thought to involve distinct sets of regions, i.e., MD and DMN regions, respectively (Fedorenko et al., 2013; Raichle et al., 2001). In contrast to this, we show the deactivation and activation of the very same set of MD regions at different junctures of difficult task episodes. Interestingly, we also found a functional distinction within the DMN regions. Bilateral temporoparietal junctions were primarily sensitive to the demand of instantiating complex programs at the beginning of hard episodes and showed differential deactivation to demands only at the beginning of episodes and not during their subsequent execution. In contrast, posterior cingulate and anteromedial prefrontal regions were only sensitive to the demand of instantiating complex control interventions and did not show greater deactivation at the beginning of difficult episodes but did show greater deactivation during their subsequent execution. While this further proves that the demands at the beginning of difficult episodes are categorically distinct from those during their subsequent execution, the reason for this functional distinction between these regions will have to await subsequent studies.

WM and attentional demands, cognitive conflict, response inhibition, rule-switching all have been seen to dilate pupils, which has been variously interpreted as reflecting increased mental load, arousal, attention, effort, etc., at junctures that involve such control demands (Strauch et al., 2022; van der Wel & van Steenbergen, 2018). Such explanations, however, cannot account for why task difficulty would cause a decrease in pupil size at the beginning of a difficult episode but an increase during its subsequent execution. For such accounts, pupil size should either have increased at the beginning of difficult episodes. A popular account has tried to explain task-related changes in pupil size in terms of exploration-exploitation trade-offs and posits that pupil size is greater during periods of exploration, allowing for increased access to the visual periphery and is smaller during exploitation when the ongoing mental task is being focused upon or ‘exploited’, which allows for more intense but narrow focus(Gilzenrat et al., 2010; Grujic et al., 2024; Jepma & Nieuwenhuis, 2011). This is based on the finding that the pupil size tends to be smaller when participants are more on-task and showing higher task performance and larger when they are less on-task and more mind wandering. While this account gives a plausible explanation for why pupil size may decrease at the beginning of difficult compared to easy episodes (‘more intense task focus or exploitation’), as per its predictions, the pupil size should have remained small on step 2 of difficult episodes because these too required a more intense task focus. None of the existing accounts predict the dichotomous response seen during hard compared to easy episodes whereby pupil size decreases at the beginning but increases subsequently. We argue that this dichotomous response results from the categorically different demands needed at the beginning of difficult task episodes when higher-level programs must be instantiated from those needed during the subsequent execution.

The evidence of programs at the beginning of diverse kinds of task episodes suggests that their role must be general and applicable to any extended task. All extended tasks require a dynamic organization of cognition across time by continually creating the correct configurational changes in various perceptual, attentional, mnemonic, and motor domains so that every instant within the task episode, cognition is anticipatorily in the most optimal state achievable for the expected demands (Bartlett & Bartlett, 1932; Logan & Gordon, 2001; G. A. Miller et al., 1960; Nobre & van Ede, 2023; Palenciano et al., 2019; Rogers & Monsell, 1995). What exactly gets done as part of these will, of course, vary across different tasks. During, e.g., a typical block or episode of visuo-motor trials, processing related to mind-wandering, ongoing unconscious goals, task-irrelevant sensory and motor processing, etc., have to be relegated (Aarts & Custers, 2012; Cardellicchio et al., 2020; Farooqui & Manly, 2018a); the predictiveness of the block needs to be utilized to make anticipatory changes, e.g., the knowledge that responses would be right-handed, visual attention limited to the area around fixation, along with an implicit idea of inter-trial intervals gets used to increase attention and make available the correct perceptual and motor processing routines at times when a stimulus is expected and decrease them when inter-trial intervals are expected, etc. (Coull & Nobre, 1998; Cravo et al., 2017; Denison et al., 2017; Ghose & Maunsell, 2002; Grabenhorst et al., 2019; Janssen & Shadlen, 2005; Los et al., 2017; Zimmermann et al., 2017). Further, cognition does not just use temporal expectations to make such anticipatory set changes but also uses them to make control interventions. Control processes are maintained across time in the way and for the duration that is expected to be most optimal based on task knowledge and past experiences. Thus, at the expected juncture, all kinds of processes get enhanced – accumulation of perceptual evidence and fine-tuning of response threshold (Fernández et al., 2019; Jepma et al., 2012), perceptual speed and acuity (Fernández et al., 2019; Vangkilde et al., 2012), control of WM (Gresch et al., 2021; Wilsch et al., 2015), arousal and cognitive resources (Shalev & Nobre, 2022; Shen & Alain, 2012), memory encoding (Jones & Ward, 2019), to list a few.

All of these occur seamlessly and automatically once the episode is embarked upon. These very many metacontrol changes through which various aspects of cognition, including control interventions, are organized across time cannot be achieved through separate and independent neurocognitive acts, and must be instantiated as part of a single goal-directed program. It is plausible that such programs correspond to configurational changes created at the beginning of the episode that adjust widespread synaptic weights in such a way that, across the ensuing task episode, neural population activity in widespread regions will get channeled through a specific sequence of states that correspond to the relevant sequence of cognitive states. Such task-related configuration of synapses will decrease ongoing but irrelevant neural activity, causing a deactivation that correlates with the length and complexity of the ensuing episode. Completion of episodes may reset these synaptic changes, corresponding to a widespread burst of activity seen at task completions (Farooqui et al., 2012; Fox, Snyder, Barch, et al., 2005; Fujii & Graybiel, 2003; Sridharan et al., 2008). In line with these, task episodes that are identical in terms of their content, e.g., rules, control demands, attentional strategies, WM etc., but differ in terms of their overarching programs because, e.g., they differ in their durations, still elicit different patterns of activity at their beginning and completions (Giray et al. in preparation).

There are limits to how big/complex a program can be instantiated or maintained at a time. Long and complex trial sequences are not executed as one task unit through a massive program; instead, they are chunked and executed as a series of smaller units, each executed through a smaller program (Farooqui et al., 2023). In comparison, short trial sequences or long but simple trial sequences get executed as a single task unit through a single program. Notably, all of this occurs even though these trial sequences did not involve WM load and hence the above cannot be attributed to WM load. The same is seen with motor sequences. As individual motor acts become easier and less demanding with practice, the length of their sequences that can be executed as a single unit through a single overarching program increases (Ramkumar et al., 2016; Tosatto et al., 2022). The same is also suggested by the nature of task labels that people employ to describe their behavior (Vallacher & Wegner, 1987, 1989). When executing easy and routine behaviors, individuals typically choose labels corresponding to long overarching tasks, e.g., saying that they are ‘preparing breakfast’ rather than ‘chopping fruit’. In contrast, when the same behavior in the same context becomes more challenging, e.g., when using a heavy and blunt knife, they become more likely to use the ‘chopping fruit’ label. When chopping fruit with a very heavy knife, fewer resources are available to maintain large programs related to long task units, and behavior gets executed as smaller task units through smaller programs.

### Summary

We showed that activation of MD regions and increased pupil size during difficult tasks is only part of the story. The beginning of these very tasks is accompanied by the deactivation of these regions and by decreased pupil size, showing that the cognitive demands of initiating difficult task episodes are categorically different from the demands related to instantiating control interventions later during these episodes. We suggested that the demands at the beginning of difficult episodes pertains of instantiating metacontrol programs that will go on to goal-directedly organize and cohere the various control interventions during the ensuing task execution.

## Materials and Methods

### MR acquisition and Preprocessing

MRI scans were acquired on a 3-Tesla Siemens MAGNETOM Tim Trio scanner with 12-channel (in Experiments 1 and 2) and 32-channel (in Experiment 3) phased array head coils. A sequential descending T2*-weighted gradient-echo planar imaging (EPI) acquisition sequence was used with the following parameters: acquisition time of 2000 ms, echo time of 30 ms, 32 oblique slices with a slice thickness of 3 mm and a 0.75-mm interslice gap, in-plane resolution of 3.0 × 3.0 mm², matrix dimensions of 64 × 64, a field of view measuring 192 mm, and a flip angle of 78°. T1-weighted MPRAGE structural images were acquired for all subjects, characterized by a slice thickness of 1.0 mm, resolution of 1.0 × 1.0 × 1.0 mm³ isometric voxels, a field of view spanning 256 mm, and 176 slices. Experiment 3 used GRAPPA and phase oversampling at 20% with an acceleration factor of 2, GRAPPA and phase oversampling were not used in experiments 1 and 2.

Preprocessing was conducted using Statistical Parametric Mapping (SPM) through automatic analysis (aa) pipeline (version 5; Cusack et al., 2015). During preprocessing, functional images were slice-time corrected using the middle slice as a reference, co-registered with the structural T1-weighted images, and followed by realignment and normalization to the Montreal Neurological Institute (MNI) template. Normalization was done through the DARTEL toolbox of SPM. EPI images were then resampled to a 2 x 2 x 2 mm³ resolution and subsequently smoothed using an 8-mm full-width at half-maximum Gaussian kernel. The dataset underwent high-pass filtering with a cut-off frequency of 128 seconds.

### Experiments 1 and 2

Participants did an auditory working memory updating task and a tactile decision-making task. The auditory working memory updating task was a modified auditory n-back task (Figure 1). On easy episodes, participants did a 1-back task, and on hard, a 3-back task. Each episode had 10 trials. On each trial, participants heard a Turkish letter. They responded if the presented letter was the same or different compared to that presented *n* trials earlier (index-finger: same, middle-finger: different). Easy and hard episodes alternated with each other. 5 to 15 seconds after the completion of an episode, participants heard rest instructions. They could rest as long as they wanted and then press a button to start the next episode. Participants completed a 10-minute pre-scan training before the fMRI session. For 3 participants whose responses remained erroneous at the end of this 10-minute session, the hard episodes were changed to 2-back.

The tactile decision-making task had participants make judgments about the size of geometric shapes (e.g., circles, squares, and rectangles) embossed on plexiglass tablets (Figure 4). Trials were organized into easy and hard episodes, each consisting of 5-trials. On each trial, participants were presented with a pair of geometric shapes, with one always being larger in size than the other. They indicated their decision using a button box. The difficulty of the decision was modulated through two main manipulations: the magnitude of the difference between areas of the two shapes and the raised-ness of the shapes’margins above the tablet.

Shapes on trials of easy episodes had a size difference between 0.8 and 1 cm^2^. On hard episodes, this difference was 0.3 to 0.5 cm^2^. Each tablet had stimuli for the 5 trials constituting an easy episode, and 5 trials constituting a hard episode. An experimenter replaced the tablet after every two episodes. The tablets were positioned either on the chest or the abdomen of participants. Participants executed 4 easy and 4 hard episodes for their pre-scan practice. During the scan, they executed 10 easy and 10 hard episodes. The experiment was coordinated through pre-recorded auditory instructions delivered through Psychtoolbox on MATLAB. These told the participants when to start a trial, when to make their button-press response, and when to rest (2 to 10 seconds after the completion of an episode). They could rest as long as they wanted and then press a button to inform that they were beginning the next episode.

### Imaging Analysis

Analysis was done using SPM. Each task episode was modeled using three regressors. The start of the episode was modeled as an event of no duration to capture the activity elicited by the beginning of the episode. The entire episode was separately modeled as an epoch of the same duration as the episode. This was to capture the average activity elicited during the episode. The completion of the episode was modeled as an event of no duration to capture the activity elicited by the completion of the episode. The easy and hard episodes were separately modeled with a different set of regressors. Movement parameters were added to the GLM as covariates of no interest. Regressors and covariates were convolved with the standard hemodynamic response function and entered into the GLM. We did two t-contrasts. The first looked at hard>easy at the beginning of the episodes. The second looked at hard>easy during the main episode epoch. Contrast estimates from each participant were entered into a group-level analysis. We then did a whole-brain repeated measures ANOVA to look for regions where the effect of these two contrasts differed.

### Experiment 3

Both easy and hard episodes started with a start-screen that stayed on until participants pressed a button and were followed by step 1 (subtask B-line presentation). The lines were presented around the center (Figure 7). The two lines presented during hard episodes were never of the same color. These lines stayed on until participants pressed a button. Immediately at the offset of these lines, a mask screen was presented for 500 milliseconds. This was followed by Step 2 (subtask A-line presentation). These, too, stayed until a button was pressed, and a mask screen also followed their offset. Participants were then probed to recall one of these A-line orientations (step 3). The probe for the line orientation to be recalled was presented at the same spot and in the same color as the original line. Participants changed the probe orientation using repeated button presses. Index and middle finger presses changed the orientation in clockwise and anticlockwise directions respectively, each press changing it by 1°. After participants were satisfied with the probe orientation, they made a separate button press to confirm their answer. This removed the probe and displayed the feedback which was the angular deviation between the actual orientation and their answer. Lastly (step 4), they recalled the orientation of one of the B lines presented in step 1. The probe again was of the same color and at the same spot as the line to be recalled. The probe manipulation was identical to that at step 3. This step, too, terminated with the confirmatory button press and was followed by the feedback that told participants of the angular deviation between their answer and the original orientation. The start-screen for the next episode followed step 4. Each of the steps was separated by a jittered gap of 0.5 to 6 seconds calculated from the offset of the previous step to the onset of the next.

Participants executed a total of 20 easy and 20 hard episodes interleaved with each other. The order re-set after every 4 episodes, e.g., for the first four episodes, the order may be easy-hard-easy-hard. For the next four, the order may shift to hard-easy-hard-easy. Data were collected through two separate fMRI runs, each run included 10 easy and 10 hard episodes and lasted between 15 to 25 minutes. An MR-safe 32-inch LCD Monitor (refresh rate: 60Hz, spatial resolution 1600 x 1200 pixels) displayed the stimuli that the participants saw through a mirror attached to the head coil. Responses were recorded using an MR-safe button box. The experiment was run using PsychoPy version 2021.2.3 (Peirce et al., 2019).

### Imaging Analysis

To capture the activity linked to each of our events of interest without making hemodynamic assumptions, we modeled 12 seconds of activity succeeding the beginning of each of the four steps using six 2-second-long finite-impulse regressors (Glover, 1999). Steps related to easy and hard episodes were separately modeled. Movement parameters were added to the GLM as covariates of no interest. We did a repeated-measures ANOVA to look for regions where the change in activity between hard and easy episodes was different across steps 1 and 2 (Figure 8).

### Experiment 4

The design of the experiment was identical to experiment 1, except that a jittered gap no longer separated the component steps; they were instead separated by a constant 1-second gap. Stimuli were delivered in a dark room on a computer monitor (NEC MultiSync LCD 2190UXP monitor, size 21 in, resolution: 1600×1200, refresh rate: 60 Hz) placed 57 cm from participants whose heads were fixed with their chins rested. Pupil sizes were recorded using Eye Tracker 6 (Applied Science Laboratories, Bedford, MA, USA) at a sampling rate of 50 Hz. Participants responded via arrow keys on a QWERTY keyboard.

The width and color of lines were chosen such that easy and hard screens were iso-luminant (https://www.w3.org/Graphics/Color/sRGB). In-house developed Python scripts were used for the analysis of pupil-size data. NaNs replaced eye-blink values. Extreme pupil sizes were discarded using a cutoff value of four standard deviations above and below the median pupil diameter for each subject. We did not apply any smoothening to the data. The timing of pupillometric and behavioral outputs was aligned via a marker sent from the experiment script to the eye-tracker via the XDAT port at the beginning. Each of the four steps of an episode ended with a button press. For each step, we took the time-series of pupil sizes over the preceding and succeeding 1 second of the button press.

### Regions of Interest

All of our fMRI results are evident in whole-brain analyses. Our main reason for doing ROI analyses was to display the pattern of activities in key MD and DMN regions. We used spherical ROIs (10 mm diameter) centered at coordinates characterized by previous studies as showing characteristics of MD and DMN regions. These were bilateral inferior frontal sulcus (IFS #41 23 29), intraparietal sulcus (IPS; #37 −56 41), anterior insula extending in the frontal operculum (AI/FO; #35 18 3), anterior pre-frontal cortex (APFC; 27 50 23 and −28 51 15). IFS, IPS, and AI were taken from Duncan (2006), and APFC was taken from Dosenbach et al (2006). DMN ROIs included: posterior cingulate cortex (PCC; −8 −56 26) and anteromedial pre-frontal cortex (aMPFC; −6 52 −2), temporoparietal junction (TPJ #54 −54 28) (Andrews-Hanna et al., 2010: Vatansever er al., 2015). All coordinates are in MNI space. ROIs were made by using the MarsbaR toolbox for SPM. ROI analyses was also done using MarsbaR. Related statistical analyses were done on JASP platform (version 0.16.4)

### Participants

Experiments 1 and 2 had 30 participants each (experiment 1: 20 females, mean age = 21.5 years, range = 18-34; experiment 2: 20 females, mean age = 20.6 years, range = 18-23 years). Participants were all right-handed and mostly undergraduates (57 undergraduates, 3 post-grads). Experiment 3 and 4 had 21 participants (13 females, 6 left-handed, mean age = 27.1 years, range = 20-43; 2 high school graduates, 14 undergraduates, 5 post-graduates). These participants did these experiments blindfolded. This is because this data was originally collected for a different study. All participants gave their written consent prior to participation. The Research Ethics Committee of Bilkent University approved all experimental protocols and procedures. All data collection occurred in National Magnetic Resonance Research Center (UMRAM), Ankara.

## Acknowledgments

This study was funded by The Scientific and Technological Research Council of Türkiye 1001 grant, numbered 120K924.

## Notes

### Competing Interest Statement

The authors have declared no competing interest.

### Summary of Updates

We revised the title of the manuscript

